# Comparison of tuning properties of gamma and high-gamma power in local field potential (LFP) versus electrocorticogram (ECoG) in visual cortex

**DOI:** 10.1101/803429

**Authors:** Agrita Dubey, Supratim Ray

## Abstract

Electrocorticogram (ECoG), obtained from macroelectrodes placed on the cortex, is typically used in drug-resistant epilepsy patients, and is increasingly being used to study cognition in humans. These studies often use power in gamma (30-70 Hz) or high-gamma (>80 Hz) ranges to make inferences about neural processing. However, while the stimulus tuning properties of gamma/high-gamma power have been well characterized in local field potential (LFP; obtained from microelectrodes), analogous characterization has not been done for ECoG. Using a hybrid array containing both micro and ECoG electrodes implanted in the primary visual cortex of two female macaques, we compared the stimulus tuning preferences of gamma/high-gamma power in LFP versus ECoG and found them to be surprisingly similar. High-gamma power, thought to index the average firing rate around the electrode, was highest for the smallest stimulus (0.3° radius), and decreased with increasing size in both LFP and ECoG, suggesting local origins of both signals. Further, gamma oscillations were similarly tuned in LFP and ECoG to stimulus orientation, contrast and spatial frequency. This tuning was significantly weaker in electroencephalogram (EEG), suggesting that ECoG is more like LFP than EEG. Overall, our results validate the use of ECoG in clinical and basic cognitive research.

## Introduction

Electrocorticography (ECoG), also known as intracranial electroencephalography (iEEG), is obtained from macroelectrodes placed subdurally on the pial surface of cortex and is widely used in drug-resistant epilepsy patients. The patients are often monitored for weeks for localization of the seizure focus, allowing (with patient’s consent) researchers to conduct cognitive and neuroscience studies^1–9^.

These studies often use power in gamma (30-70 Hz) and high-gamma (>80 Hz) ranges to make inferences about the underlying neural processing^10^. High-gamma activity (>80 Hz) refers to power over a broad range of frequencies above the gamma band that, in ECoG, is modulated by stimulus presentation as well as the behavioral state^4, 5, 10–13^. High-gamma activity is also observed in local field potential (LFP) obtained by inserting microelectrodes in the cortex of animals, where it is tightly correlated with the spiking activity of neurons in the vicinity of the microelectrode^13–17^.

Gamma rhythm (30-70 Hz), which is different from high-gamma activity^17^, has been extensively studied in electroencephalogram (EEG) in humans and LFP in animals, and has been associated with high level cognitive functions such as attention, memory and perception^18–24^. Further, gamma is known to be strongly induced by stimuli such as bars/gratings and depends on stimulus properties such as size, orientation, spatial frequency, contrast and temporal frequency^16, 17, 25–29^. Stimulus dependence of gamma has also been characterized in EEG/MEG studies^30–35^. However, only a few studies have characterized the stimulus preference of gamma in ECoG^30, 36^. No study, to our knowledge, has done a direct comparison of stimulus preferences of gamma/high-gamma in LFP versus ECoG.

Apart from providing clues about the neural correlates of gamma/high-gamma activity in ECoG, such a comparison allows us to determine the spatial spread (the cortical area around the electrode that contributes to the signal that is recorded from that electrode) of ECoG, which we have recently shown to be very local^37^. For example, both the firing rates and LFP high-gamma power reduce with increasing stimulus size because of larger surround suppression^17^. However, since a larger stimulus activates a larger cortical area, we might observe an increase in ECoG high-gamma (despite a reduction in firing rate) if ECoG spatial spread is much larger than LFP. Similarly, we have recently shown that gamma power recorded using EEG has much weaker tuning preferences (for stimulus orientation, size and contrast) compared to LFP^29^. A comparison of analogous gamma tuning preferences for ECoG versus LFP will provide clues about their similarity. Recording from a unique hybrid grid which consists of both micro and macro-electrodes, implanted in the primary visual cortex of the same two female macaques for which we had earlier compared LFP versus EEG tuning^29^ and LFP versus ECoG spatial spreads^37^, we compared the strength of ECoG and LFP gamma/high-gamma power for different stimulus properties such as size, orientation, spatial frequency and contrast.

## Results

We simultaneously recorded LFP and ECoG signals using a special custom-made hybrid grid electrode array implanted in the left primary visual cortex (V1) of two monkeys (Monkeys 3 and 4), trained to perform a fixation task, while visual gratings that varied in size, orientation, contrast or spatial frequency were presented on a screen. This hybrid grid consisted of 9 (3×3) ECoG electrodes and 81 (9×9) microelectrodes, both attached to the same connector and referenced to same wire. The microelectrode array was placed between four ECoG electrodes in V1 (see Figure 1 of Ref 37). For the variable stimulus size condition (Figures 1-4), data from two additional monkeys (Monkeys 1 and 2) was used, for which microelectrode and ECoG recordings were conducted separately (see Methods for details). All spectral analyses were performed using the multi-taper method^38, 39^.

**Figure 1:**
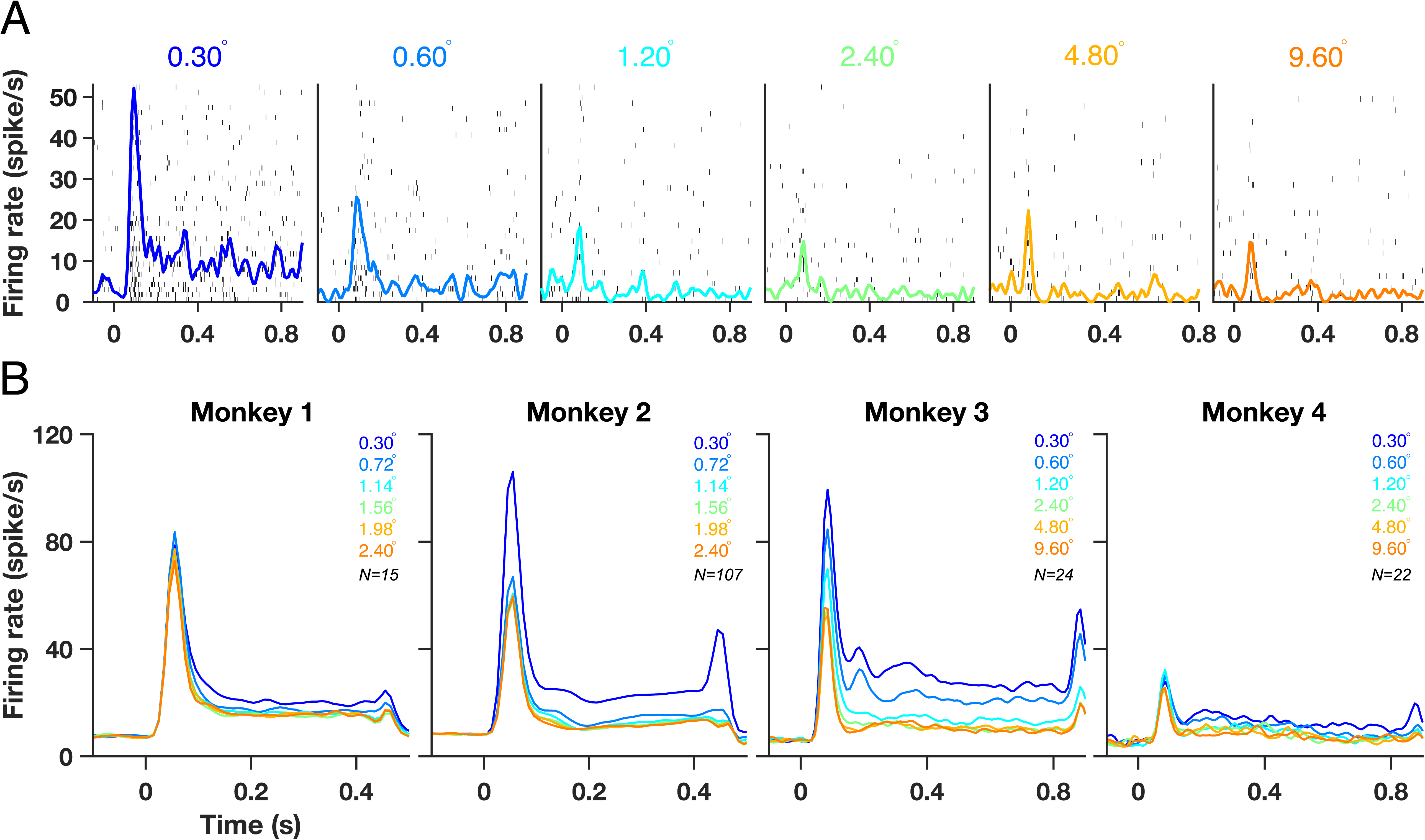
Spiking activity for different stimulus sizes. **(A)** Raster plots showing spiking activity in individual trials for each stimulus size for an example unit from Monkey 3. Each row represents a trial. The peristimulus histogram, averaged across trials is overlaid on the raster plots. **(B)** Averaged firing rates for six stimulus sizes shown as different color traces for Monkeys 1, 2, 3 and 4.

### High-gamma activity in ECoG is maximum for a small stimulus size (radius of 0.3°)

Figure 1A shows the raster plot and multiunit firing rate of an example recording site from Monkey 3 when gratings of six different radii (0.3°, 0.6°, 1.2°, 2.4°, 4.8° and 9.6°) were presented between 0 and 800 ms. The peristimulus histogram averaged across trials is overlaid on each of the raster plots. Consistent with our previous results^17^, increasing the stimulus size decreased the firing rate. Similar trends were observed for the population dataset of 15, 107, 24 and 22 recordings sites from the four monkeys (Figure 1B). Note that the stimulus radii for Monkeys 1 and 2 were different from Monkeys 3 and 4.

Next, we studied the LFP and ECoG signals for varying stimulus sizes. Figure 2A shows the change in LFP power relative to the baseline period (defined as 500 to 0 ms before stimulus onset) for the same example site as Figure 1A from Monkey 3 for six different sizes. These time-frequency energy difference spectra showed a prominent gamma rhythm (red horizontal band) at ∼50 Hz for stimulus size of 0.6° and above, which appeared after the initial transient and remained present throughout the stimulus duration (up to 0.8 s). Consistent with previous studies^16, 17, 26, 29^, strength of LFP gamma rhythm increased with an increase in stimulus size while the gamma peak frequency decreased. Further, the smallest stimulus (radius 0.3°) showed a prominent increase in power over a broad frequency range above the gamma band. The power in this broadband showed the opposite trend and decreased with an increase in stimulus size. Figure 2B shows the time-frequency difference spectra for an example ECoG electrode from the same monkey. Similar to LFP, the power of ECoG gamma increased with increasing stimulus size. Surprisingly, even though the ECoG electrode was much larger than LFP, the smallest stimulus produced the largest high-gamma power even in ECoG. The increase in ECoG high-gamma power was more prominent up to ∼250 Hz, unlike LFP high-gamma that remained prominent up to 400 Hz and beyond. Similar results were obtained from the population average of 24 LFP sites and 5 ECoG sites (Figure 2C and 2D).

**Figure 2:**
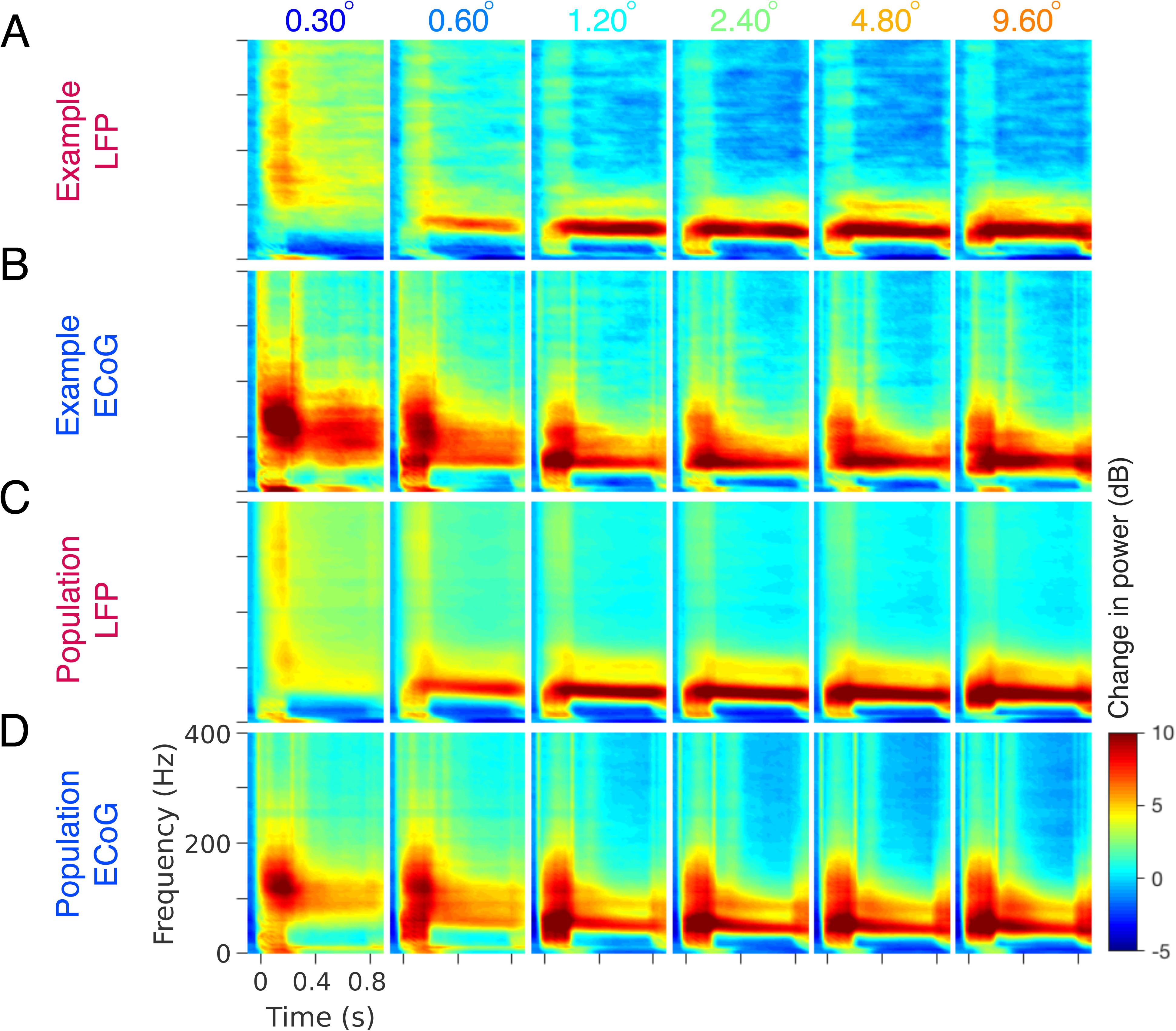
Gamma oscillations and high-gamma activity as a function of stimulus size in LFP and ECoG for Monkey 3. **(A)** Time-frequency energy difference plots (in dB) showing the difference in energy relative to baseline energy (−500 to 0 ms, 0 denotes the stimulus onset, stimulus is presented from 0 to 800 ms) for six stimulus radii (labelled above the plots in degrees) for an example LFP recording site (same as shown in Figure 1A). The gamma rhythm at ∼50 Hz increases with size, while the high-band activity above the gamma band decreases with size. **(B)** same as A for an example ECoG recording site. **(C–D)** show the corresponding population responses of 24 LFP and 5 ECoG recording sites.

Figure 3 A, C, E and G show mean change in power from the baseline (obtained by subtracting log of baseline power from the log of stimulus power, see Methods for details) across recording sites, as a function of frequency for Monkeys 1, 2, 3 and 4. In all monkeys, the largest stimulus produced the strongest but slowest gamma, visible as a prominent peak at ∼45-60 Hz (orange traces). In all monkeys except Monkey 4, a prominent harmonic of gamma was also visible between 80-120 Hz. However, there were interesting differences between Monkeys 1, 2 and Monkeys 3, 4, because much larger stimulus sizes were used for the latter two monkeys. For example, in Monkey 4, a second gamma peak was clearly visible at ∼30 Hz for the largest stimulus size, which is the ‘slow’ gamma as described in our previous study^29^. Also, the LFP gamma in Monkey 4 was weaker than Monkey 3 (this was also observed in our previous study^29^, in which recordings were done from a different hemisphere using a different array); we discuss this in more detail in the Discussion. Importantly, in spite of the differences in the strength of gamma and high-gamma band across monkeys, the overall trends remained similar: the strength of gamma rhythm increased with an increase in stimulus size whereas high-gamma power decreased. Importantly, similar trends were also observed in the ECoG signals. To compare the changes in power with stimulus size for LFP and ECoG, we computed the power in two frequency bands: 30-65 Hz for gamma and 150-250 Hz for high-gamma, as shown in Figure 3 B, D, F and H. The gamma range was chosen to avoid the ‘slow’ gamma, while the high-gamma range was chosen to avoid the harmonic of gamma between 80-120 Hz. As observed in PSD plots, the power in gamma band increased with size for both LFP and ECoG (the only exception was the ECoG of Monkey 2 for which only a single electrode was available), whereas high-gamma power showed opposite trends. Interestingly, high-gamma power was maximum for the smallest stimulus (radius of 0.3°) for both LFP and ECoG for all the four monkeys. This suggests local origins of ECoG in primary visual cortex, similar to our previous study^37^, since high-gamma would have been expected to be higher for a larger stimulus if spatial summation occurred over a large cortical area for ECoG. However, unlike our previous approaches^37^, this approach did not provide a quantitative estimate of the spatial spread. We discuss this in more detail in the Discussion.

**Figure 3:**
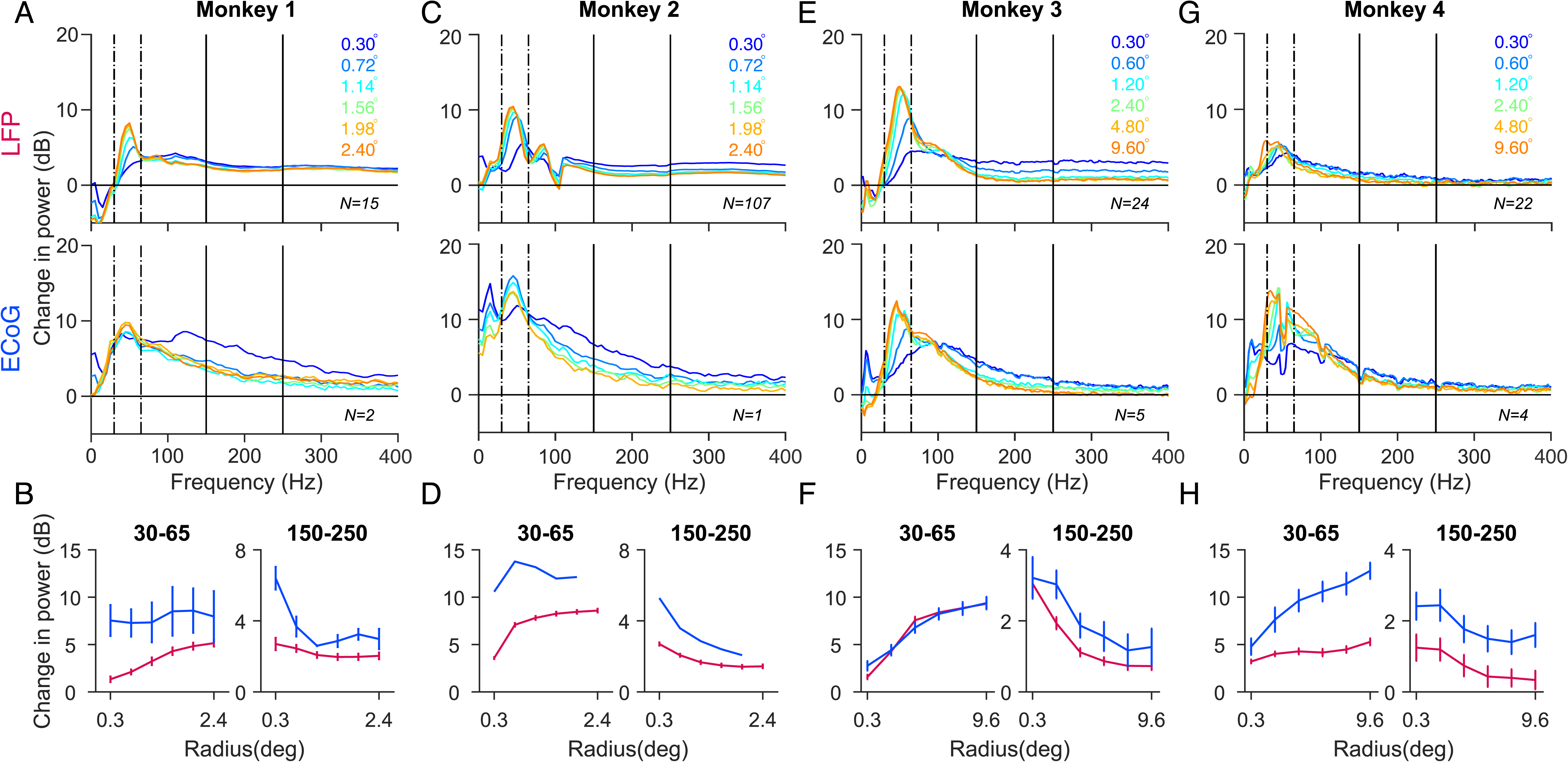
Tuning of gamma oscillations and high-gamma activity for stimulus size. **(A, C)** Average relative change in power spectra between 200 and 400 ms from baseline energy (−200 to 0 ms) for 15 and 107 LFP recordings sites (top panel), 2 and 1 ECoG recording sites (bottom panel) for Monkeys 1 and 2. **(E, G)** same as A, C but for 24 and 22 LFP recordings sites (top panel), 5 and 4 ECoG recording sites (bottom panel) for Monkeys 3 and 4. The change in power is computed between 250 to 750 ms relative to baseline energy (−500 to 0 ms). **(B, D, F and H)** Change in LFP (magenta) and ECoG (blue) for gamma (30 – 65 Hz) and high-gamma (150 - 250 Hz) frequency bands as a function of stimulus size. Error bar indicates SEs of the mean. Note that the stimulus radii for Monkeys 1 and 2 are different from Monkeys 3 and 4.

**Figure 4:**
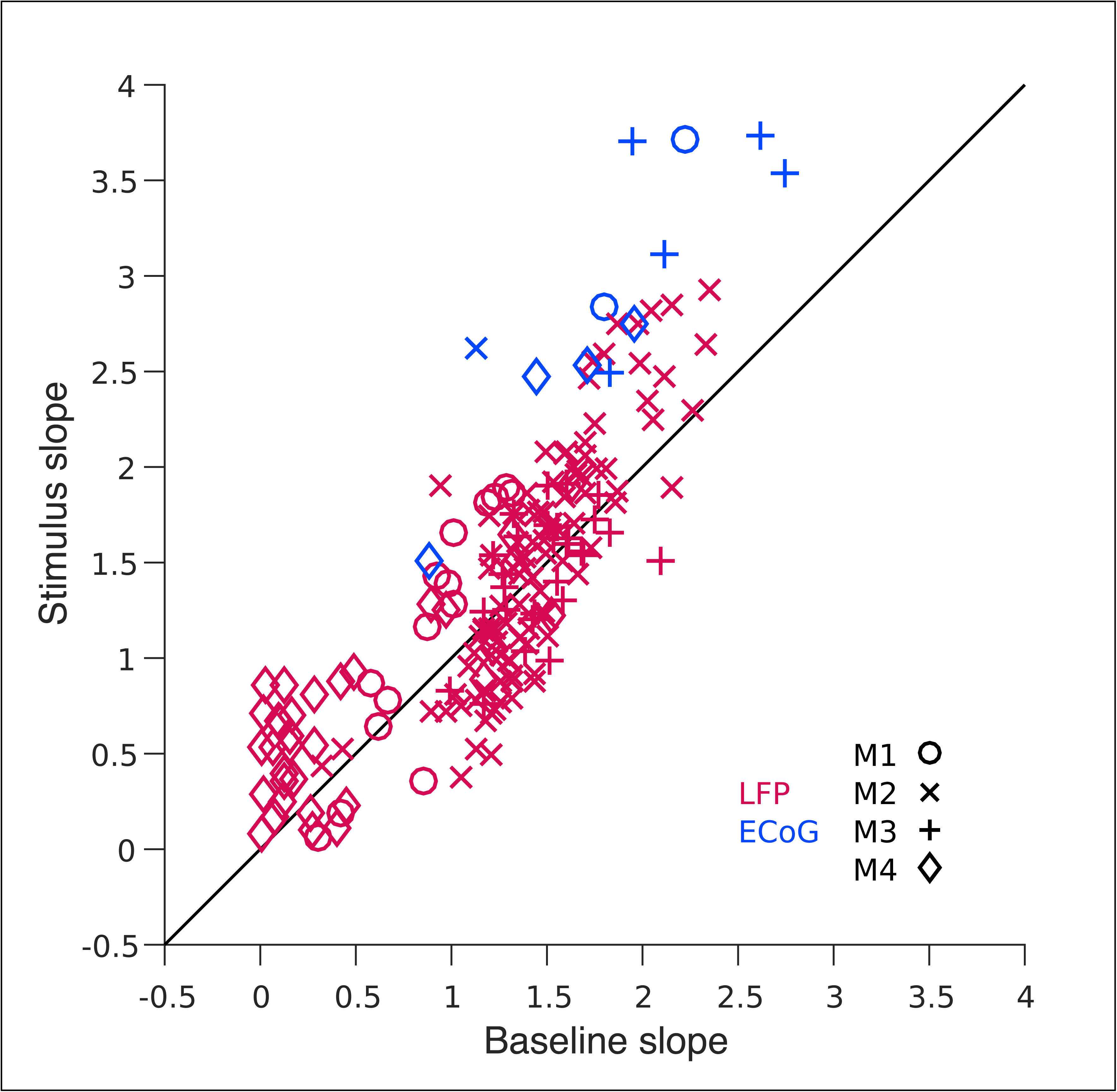
Slope of the high-gamma activity for 0.3° stimulus. The slope of LFP (magenta) and ECoG (blue) electrodes computed for high-gamma frequency range (150 – 250 Hz) for baseline period is plotted in x-axis and for stimulus period in y-axis. The four monkeys are represented using four different marker types.

A comparison of the shape of the change in power spectra for LFP (Figure 3A, C, E, G, top row) versus ECoG (bottom row) revealed an interesting difference. Beyond ∼100 Hz, the traces were almost parallel to the x-axis in the case of LFP (in all except Monkey 4) but showed a negative slope for ECoG in all monkeys. This suggested that the slope of the PSD in the high-gamma range during stimulus and baseline periods were comparable in case of LFP (such that the difference produced a zero-slope line), but stimulus PSD had a steeper slope than baseline in case of ECoG. Indeed, we have previously observed that while increase in high-gamma power could be observed up to at least ∼400 Hz in LFP^40^, it was prominent only up to ∼150 Hz in human ECoG^13^. We further quantified this by plotting the slopes of high-gamma range during stimulus period versus baseline (Figure 4). The LFP slopes for stimulus and baseline period were comparable (mean slope during stimulus: 1.31, baseline: 1.22, p=0.15, paired t-test (two sample t-test)), whereas the ECoG slopes for stimulus period were greater than baseline period (mean slope during stimulus: 2.92, baseline: 1.87, p=0.00035).

### Stimulus tuning of gamma oscillations

We first compared the orientation tuning (both preferred angle and selectivity; equations 3 and 4) between LFP and ECoG, for two reasons. First, while it is well established that different neurons prefer different orientations in V1 such that the distribution of orientation preferences of MUA is more or less uniform^41–43^, several studies have shown that the stimulus orientation that generates the strongest gamma in microelectrode recordings is remarkably similar across all the recording sites^16, 29, 44^. However, since these microelectrode arrays span only ∼4×4 mm^2^ patch of cortex, it is possible that different patches of cortex prefer different orientations (the preferred orientation for gamma is location specific, but not monkey specific). Because ECoGs record from brain areas separated by 10 mm or more, comparison of orientation preferences across ECoG sites could provide clues about the specificity of orientation tuning in the gamma band. Second, we have recently shown that the orientation selectivity (measure of the strength of orientation tuning) for gamma was much weaker in EEG compared to LFP^29^. This could be because EEG records activity from a much larger part of the brain than LFP, and these parts may not be as well tuned for a particular orientation. A comparison of the orientation selectivity of ECoG and LFP could therefore provide clues about their similarity.

Figure 5A shows the population average of the change in LFP and ECoG power as a function of frequency, across 77 LFP (top) and 5 ECoG (bottom) recording sites for Monkey 3. The change in power was computed between 250 ms to 750 ms relative to baseline period (0 ms to 500 ms before stimulus onset) and then averaged across sites on a log scale. The eight colored traces represent the change in power spectrum for eight stimulus orientations. We observed that the mean LFP gamma between 45 to 70 Hz was strongest and fastest at a stimulus orientation of 90°. Surprisingly, mean ECoG gamma showed similar trends as LFP gamma with the strongest and fastest gamma for 90° orientation (Figure 5B, top panel).

**Figure 5:**
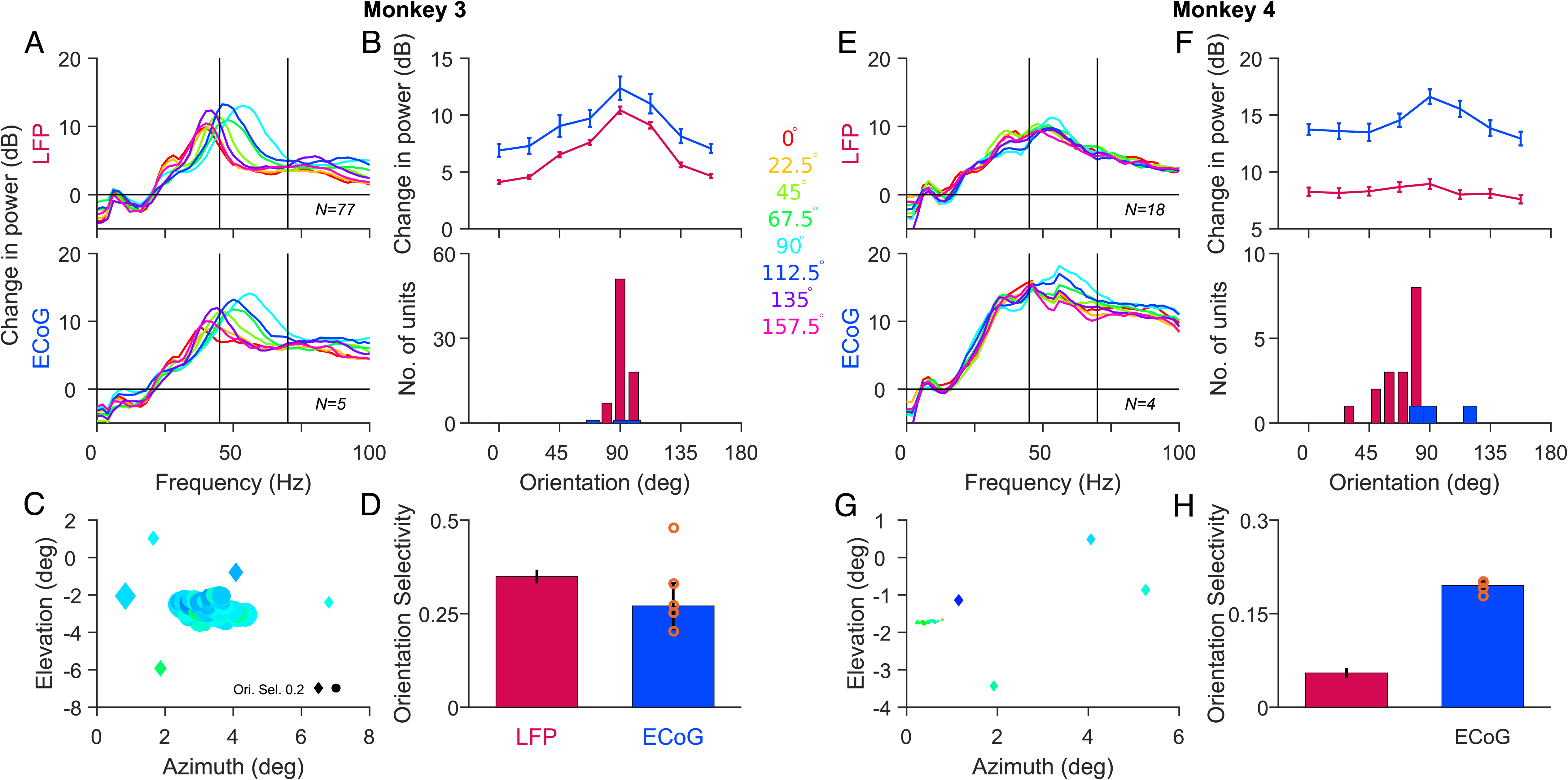
Orientation tuning of gamma oscillations in LFP and ECoG. **(A)** Average relative change in power spectra between 250 and 750 ms from baseline energy (−500 to 0 ms) for 77 LFP (top panel) and 5 ECoG recording sites (bottom panel) for Monkey 3. Eight colored traces are for eight different orientation values (labelled at the centre of Figure). **(B)** Average change in gamma power as a function of orientation (top panel) and the histogram of orientation preference (bottom panel) across recording sites for LFP (magenta) and ECoG (blue). Error bar indicates SEs of the mean. **(C)** Orientation preference of gamma rhythm across LFP (circle) and ECoG (diamond) recording sites plotted at the respective RF centers. The color represents the preferred orientation while the size of the marker represents the strength of tuning. **(D)** Median orientation selectivity of LFP and ECoG across recording sites. Error bar indicates SEs of the median, computed using bootstrapping. The orange circles are the five ECoG electrodes. **(E–H)** same as **A–D** but for 18 LFP and 4 ECoG recording sites in Monkey 4.

To examine the preferred orientation of gamma at different cortical locations we computed the preferred orientation of gamma in 45 to 70 Hz frequency range for each of the recording sites. Figure 5C shows ECoG (diamonds) and LFP (circles) electrodes, plotted at their receptive field centers and color-coded based on preferred orientation for Monkey 3. Consistent to previous studies^16, 29, 44^, we observed that preferred orientation of LFP gamma was similar across sites (Figure 5B, bottom panel, magenta bars). Interestingly, all the five ECoG electrodes which covered ∼20 × 20 mm in the cortex, showed a remarkably similar preference for stimulus orientation. Although we observed small variations in preferred orientation from the electrode to electrode, the distribution of ECoG (ranging from 70° to 100°) was similar to the LFP (ranging from 80° to 100°; Figure 5B, bottom panel). Further, the strength of orientation tuning measured by orientation selectivity was on average comparable for ECoG and LFP (Figure 5D). The ECoG electrodes which showed a deviation from 90° had low orientation selectivity values, represented by the smaller marker size in Figure 5C. Similar results were observed for Monkey 4 across 18 LFP and 4 ECoG recording sites (Figure 5E - 5H). Thus, the orientation preference of gamma is monkey specific but not location specific.

The orientation preference and selectivity depended on the choice of the frequency band. In particular, for Monkey 3, gamma peak frequency was below our lower cutoff of 45 Hz for some orientations. We used this gamma range to be in congruence with our previous study^29^, in which we had recorded from the same monkeys but used a microelectrode array implanted in the other hemisphere, and had also collected simultaneous EEG data. Since the orientation preferences for LFPs were similar for the two arrays, having the same frequency range allowed us to better compare the LFP, ECoG and EEG gamma tuning. Further, the low frequency cutoff could not be lowered due to the presence of ‘slow gamma’ (see Ref 29), which peaked between 30-35 Hz for the two monkeys. As discussed in more detail later, tuning properties critically depend on the choice of the lower frequency cutoff. Nonetheless, visual inspection of Figures 5A and 5E reveals that the gamma peaks were remarkably similar for LFP and ECoG for both monkeys, such that choosing a different frequency range changed the tuning parameters in similar ways.

Like orientation, gamma tuning of LFP and ECoG were similar for spatial frequency (Figure 6) and contrast (Figure 7). In particular, ECoG gamma peak frequency increased with contrast and was similar to LFP peak frequency in both monkeys (for contrasts above 25% that generated salient gamma peaks; Figure 7B, D), unlike EEG gamma peak frequency that did not show a substantial increase with contrast^29^. Overall, our results suggest that ECoG is more similar to LFP than EEG.

**Figure 6:**
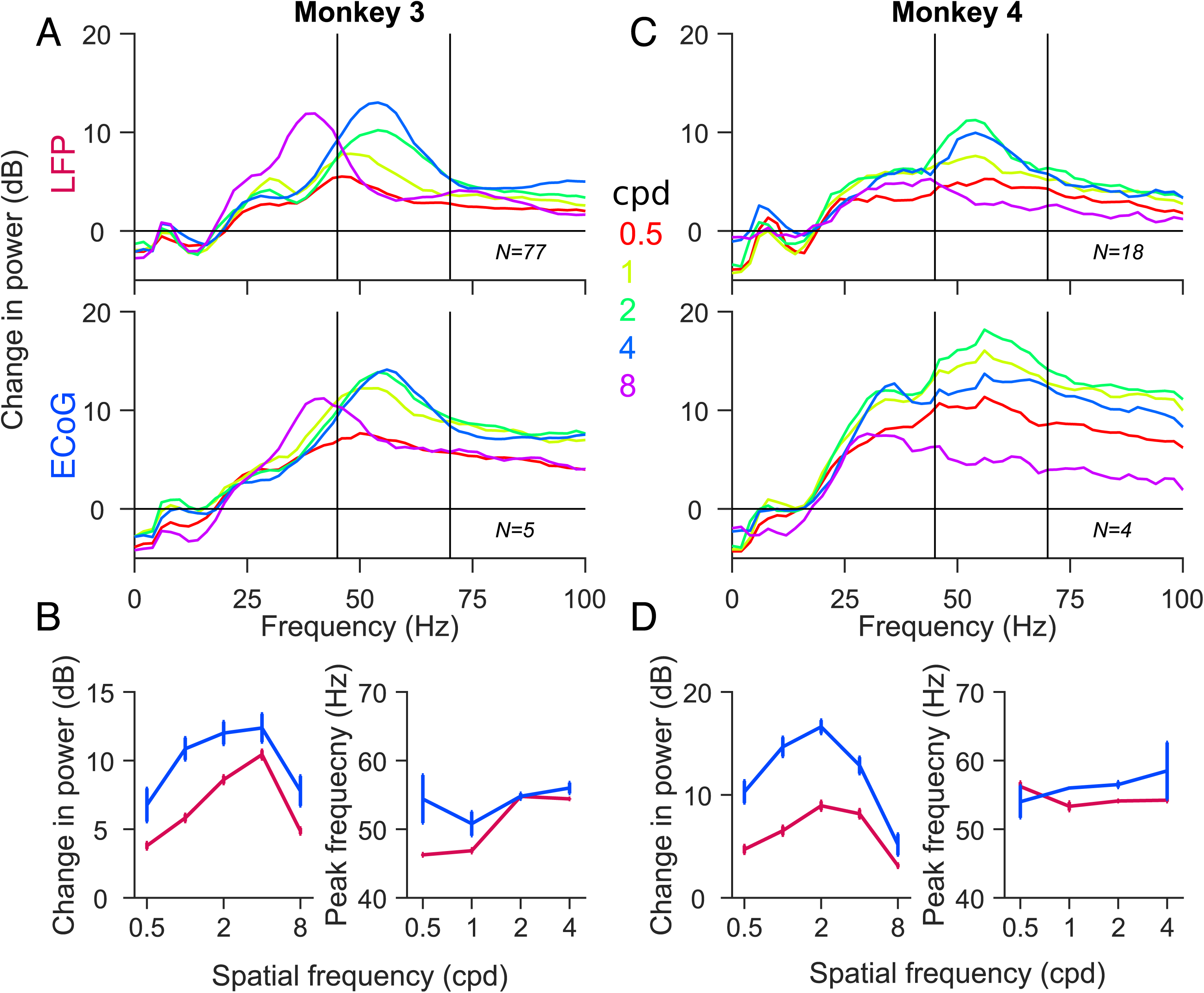
Spatial frequency tuning of gamma oscillations in LFP and ECoG. **(A, C)** Mean change in power spectra across 77 and 18 LFP recording sites (top panel), 5 and 4 ECoG recording sites (bottom panel) for Monkeys 3 and 4 calculated at stimulus orientationss that induce largest power change in gamma (90° for both monkeys). Five colored traces represent five different spatial frequency values. **(B, D)** left panel: Average change in gamma power as a function of spatial frequency for LFP (magenta) and ECoG (blue). right panel: Average gamma peak frequency as a function of spatial frequency. 8 cpd was ignored as the gamma peak was out of the selected frequency range. Error bar indicates SEs of the mean.

**Figure 7:**
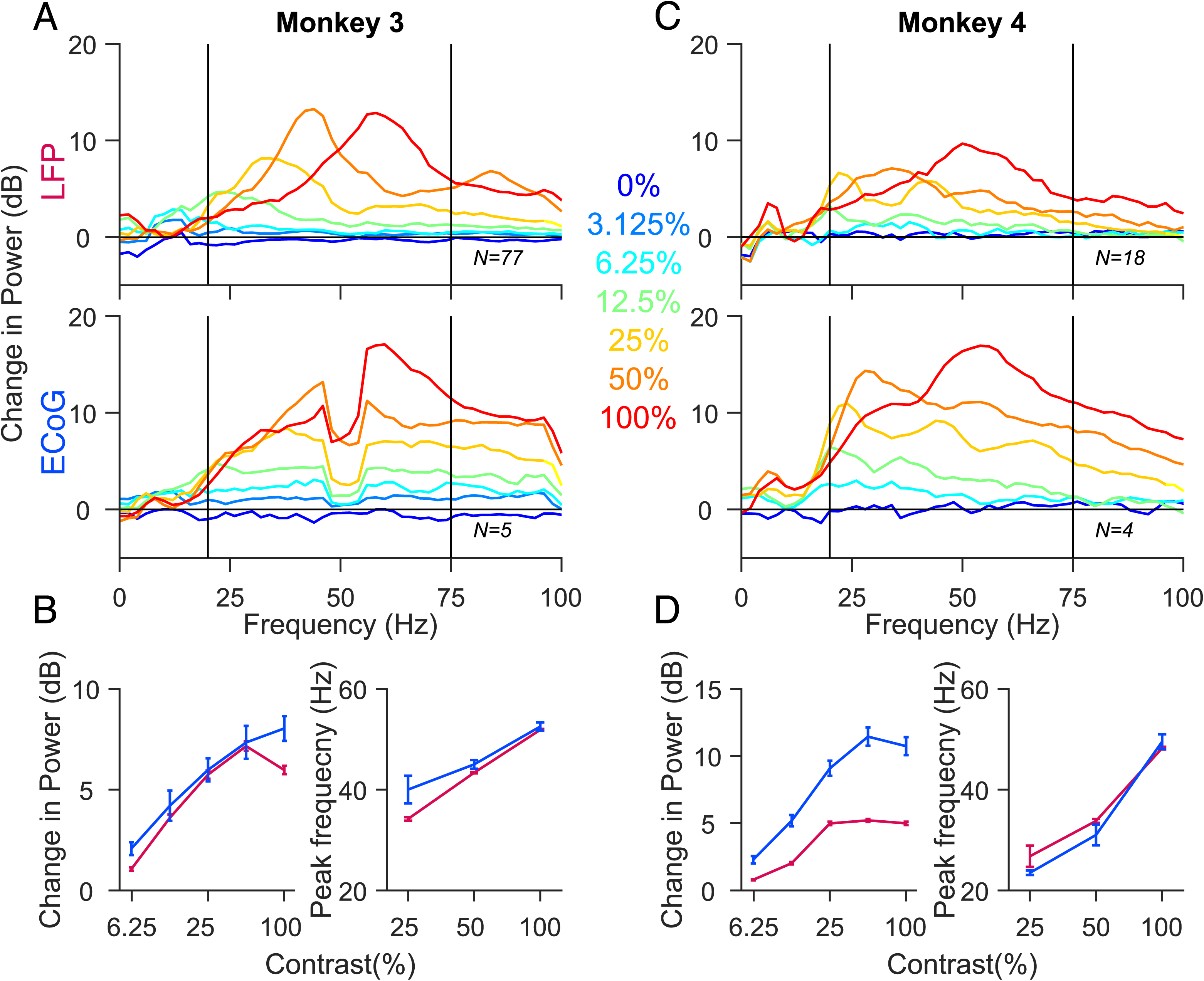
Contrast tuning of gamma oscillations in LFP and ECoG. **(A, C)** Mean change in power spectra across 77 and 18 LFP recording sites (top panel), 5 and 4 ECoG recording sites (bottom panel) for Monkeys 3 and 4 calculated at stimulus orientations and spatial frequencies that induce largest power change in gamma (90° and 4cpd for Monkey 3 and 90° and 2cpd for Monkey 4). Seven colored traces represent seven different contrast values. Note that for Monkey 4 there are only six traces. **(B, D)** left panel: Average change in gamma power as a function of contrast for LFP (magenta) and ECoG (blue). right panel: Average gamma peak frequency as a function of contrast. Error bar indicates SEs of the mean.

## Discussion

We compared the stimulus tuning properties of gamma/high-gamma in LFP and ECoG by simultaneously recording these signals using a custom-made hybrid grid and found them to be surprisingly similar. The smallest stimulus size tested (radius of 0.3°), which has been earlier shown to produce largest high-gamma power in LFP^17^, produced the largest high-gamma power in ECoG as well. Further, tuning preferences of gamma oscillations for stimulus size, orientation, spatial frequency and contrast were very similar for LFP and ECoG. Overall, these results suggest that ECoG is an excellent signal to study gamma oscillations.

These results are consistent with our recent study^37^, in which we used a receptive field (RF) mapping approach to show that the spatial spread of ECoG was surprisingly local (SD of ∼1.5 mm or 2SD of ∼3mm), not much larger than the diameter of the ECoG electrode (2.3 mm), and only ∼3 times the spread of LFP (2SD of ∼ 1mm). These results are also consistent with the observation that the RFs of ECoGs recorded in humans are very small^3^, although in that study the RFs (measured in degrees) were not converted to cortical spreads (measured in mm).

Unfortunately, this approach did not yield a quantitative estimate of the ECoG spread, for two reasons. First, it is possible that ECoG preferentially samples neurons in the upper layers of the cortex that may prefer smaller stimulus sizes, so it is difficult to deduce spatial spread from size tuning. Second, the range of stimulus sizes that we used was not wide enough to quantitatively compare the spreads of LFP and ECoG. Use of even smaller stimuli (for example, radius of 0.1°) would have yielded a better estimate of the ‘optimal’ stimulus size for LFP high-gamma power, and comparison of optimal stimulus sizes for LFP and ECoG would have yielded a quantitative estimate of their respective spatial spreads. However, when extremely small stimuli are used, appropriate comparison is possible only in the absence of eye jitters. Given that the monkey had to maintain fixation only within 1° or more around the fixation spot, it is possible that a very small stimulus would occasionally miss the receptive field completely if the monkey’s gaze was away from the fixation spot, increasing the variability of the estimate of high-gamma power for very small stimuli. The method used in our previous study^37^, which is originally based on the model proposed by Xing and colleagues^45^, partially addressed this concern because the inflation in the estimate of the RF size due to several factors (including eye jitters) is similar for different measures (MUA, LFP and ECoG), and therefore a model that estimates the spatial spreads based on the differences in RF sizes between measures (such as MUA versus LFP and LFP versus ECoG) can cancel out these common terms (see Refs ^37, 45^ for details). We had also used another approach that involved the comparison of the PSDs of ECoG and LFP during spontaneous periods to show that the ECoG spread was local. The present approach, obtained by simply comparing the high-gamma power as a function of stimulus size, provides a third, albeit weaker line of evidence that ECoG is a local signal. Further, this result is obtained without any model or additional assumptions and is complementary to the previous two approaches that used either very small stimuli to map RFs or compared the PSDs during spontaneous periods.

What are the origins of high-gamma activity in ECoG? High-gamma activity was initially interpreted in the same conceptual framework as gamma oscillations, just operating at a higher frequency^46–48^. More recently, high-gamma in the LFP has been shown to be tightly correlated with the multiunit firing rate^13–17^. ECoG high-gamma power has been proposed to reflect the synchrony in neural population^13^, although direct experimental evidence, to our knowledge, is lacking. In the size study, we observed that upper range of ECoG high-gamma was limited to 200-250 Hz compared to at least 400 Hz in LFP (see Figure 2B vs 2A for stimulus radius of 0.3°). This was consistent across electrodes (Figure 2D vs 2C) and monkeys (Figure 3A, C, E and G; bottom vs top panel), and was further quantified by comparing the slopes in stimulus period with baseline period (Figure 4). This could be because the PSD of the ECoG was much steeper than LFP at low frequencies (see Ref ^37^), and therefore the overall power of the ECoG at high frequencies was much lower than LFP. Thus, the noise (either in the device or the brain) could have affected the ECoG signal more than LFP at high frequencies. It appears that even the LFP for Monkey 4 was more affected by noise, since the PSD slopes in this monkey were shallower during both baseline and stimulus periods compared to other monkeys (Figure 4). The differences in PSD slopes for ECoG compared to LFP could be due to its larger size, lower impedance or position.

We observed that the tuning preferences of gamma were similar for ECoG and LFP for all the four stimulus manipulations (size, orientation, spatial frequency and contrast), while previously we had observed considerable differences between LFP and EEG tuning^29^. Note that while these recordings were done on the same monkeys, we did not record all three signals simultaneously because of technical difficulties (see Methods). Nonetheless, the weak tuning of EEG gamma was observed in humans also^29^, and is therefore likely to be a general feature of EEG signals. However, note that the similarity in tuning profile of LFP and ECoG gamma rhythms for different stimulus manipulations could be because of a coherent network because of the use of full screen gratings at full contrast which are known to produce strong and coherent gamma rhythms^16, 17^ over a large brain area. Both the microelectrodes and macroelectrodes captured the activity of this network and therefore showed similar tuning preferences. Interestingly, ECoG electrodes which were on the surface of cortex captured this activity as reliably as microelectrodes which were presumably in the superficial layers of the cortex. Apart from the stimulus, another factor that could have influenced our results is volume conduction^49, 50^. In a previous study^50^, in which we recorded from microelectrodes implanted in Monkeys 1 and 2, we showed that the LFP-LFP phase coherence almost becomes flat for CSD (current source density, a double spatial derivative of potential, obtained by subtracting the potential of an electrode from the potentials of four neighboring electrodes; see Fig 4A of Ref ^50^). Since, we had only five (Monkey 3) and four (Monkey 4) ECoG electrodes, the CSD analysis could not be performed for ECoG in the current setup.

As described earlier in Results section, the tuning parameters depended critically on the low frequency limit of the gamma band. This is because the actual power (not change in power which is displayed in the figures) falls off rapidly with frequency and displays a prominent “1/f” structure. The total power in a band is therefore dominated by the lower frequencies that have larger absolute power. For example, in the orientation tuning experiment, gamma peak was strongest for the stimulus orientation of 90° but also the fastest (peak around ∼55 Hz) for Monkey 3 (Figure 5A). Orientation of 0° produced a smaller bump, but since it was around 40 Hz, the power between 35-40 Hz was more for 0° stimulus than 90°. However, if we had chosen the gamma band between 35-70 Hz, the preferred orientation would have shifted towards 0° just because the absolute power between 35-40 Hz far exceeds the power between 50-60 Hz. This issue can be partially addressed by using the normalized instead of absolute power while computing the power in a band, but in general, it is difficult to compare gamma power across stimulus conditions when the peak frequency itself shifts with stimulus.

In our case, the choice of frequency band is of less relevance because the actual power spectra for LFP and ECoG were remarkably similar for every stimulus condition: if the gamma peak did not fall in a specified range for LFP, it invariably fell outside the range for ECoG as well. Therefore, our main result that LFP and ECoG gamma tuning is remarkably similar holds irrespective of the choice of the frequency band.

Although the overall trends were similar for Monkeys 3 and 4, the strength of tuning was different. For example, orientation selectivity was different for the two monkeys for LFP gamma whereas ECoG gamma showed comparable selectivity (Figure 5D and 5H). One reason could be because the LFP receptive field locations were very foveal in case of Monkey 4 (Figure 5G), although the foveal ECoG electrodes in both the monkeys showed strong orientation tuning (Figure 5C and 5G). Moreover, Xu and colleagues ^51^ found no difference in orientation selectivity as a function of eccentricity in V1. We suspect that the main reason behind weaker LFP gamma in Monkey 4 is because the microelectrode array had earlier been explanted (see Methods for details), although it is unlikely that this affected any of the major results.

To conclude, our findings highlight the presence of gamma oscillations in ECoG which shows similar tuning preference to gamma oscillations observed in LFP recordings, even though the size of the ECoG electrode is several hundred times larger than the microelectrode. Therefore, ECoG gamma can act as a potent marker for the diagnosis of brain disorders such as autism and schizophrenia which have been associated with abnormal gamma rhythms^52, 53^. Further, comparing the high-gamma activity between ECoG and LFP we showed that ECoG has local origins in V1. Together, our results validate the use of ECoG in brain-machine interface applications and basic science research.

## Methods

### Animal preparation and Recording

All animal experiments and protocols performed in this study are in strict accordance with the relevant guidelines and regulations approved by the Institutional Animal Care and Use Committee of Harvard Medical School (for Monkeys 1, 2) and Institutional Animal Ethics Committee (IAEC) of the Indian Institute of Science and the Committee for the Purpose of Control and Supervision of Experiments on Animals (CPCSEA) (for Monkeys 3 and 4). The details of our experiment design and data collection have been described in detail in our previous study^37^; here we explain them briefly. The microelectrode and ECoG data used in this study were collected in two separate set of experiments. The first set was conducted on two male monkeys (*Macaca mulatta*; 11 and 14 Kg); animal protocols approved by the Institutional Animal Care and Use Committee of Harvard Medical School. For this set of experiments, microelectrode and ECoG recordings were performed separately and are described in detail elsewhere ^17, 27, 54^. Briefly, after monkeys learned the behavioral task, a 10×10 microelectrode grid (96 active channels, Blackrock Microsystems) was implanted in the right primary visual cortex (∼15 mm anterior to the occipital ridge and ∼15 mm lateral to the midline). The microelectrodes were 1 mm long separated by 400 μm. After microelectrode recordings, a second surgery was performed to implant the custom-made array having 2 ECoG contacts (2.3 mm in diameter and 10 mm apart, Ad-Tech Medical Instrument) on the left primary visual cortex of the same monkeys (see Materials and Methods of Ref 37, for details). One ECoG electrode in Monkey 2 did not show any stimulus evoked response and thus was excluded, yielding 3 ECoG electrodes from these two monkeys. Note that ECoG and microelectrode recordings were non-simultaneous for these two monkeys.

The second set of experiments involved simultaneous recordings of spikes, LFP and ECoG signals from two female adult monkeys (*Macaca radiata*; 3.3 and 4 Kg); animal protocols approved by the Institutional Animal Ethics Committee (IAEC) of the Indian Institute of Science and the Committee for the Purpose of Control and Supervision of Experiments on Animals (CPCSEA). Once the monkey had learned the fixation task, a custom-made hybrid array (see Figure 1 of Ref 37) was implanted in the left cerebral hemispheres. This hybrid array had 3×3 ECoG electrodes (Ad-tech Medical Instrument) and 9×9 microelectrodes, both attached to the same connector made by Blackrock Microsystems. The ECoG electrodes were platinum discs of exposed diameter of 2.3 mm and inter-electrode center-to-center distance of 10 mm. The microelectrodes were 1 mm long, 400 μm apart. The electrode array was implanted under general anesthesia; first a large craniotomy and a smaller durotomy were performed, subsequent to which the ECoG sheet was inserted subdurally such that the previously made silastic gap between four ECoG electrodes was in alignment with the durotomy (see Ref 37 for details). The microelectrode array was finally inserted into the gap, ∼10 – 15 mm from the occipital ridge and ∼10-15 mm from the midline. In Monkey 3, out of six ECoG electrodes which were posterior to lunate sulcus, one had noisy receptive field estimate, yielding 5 ECoG electrodes for further analysis. For Monkey 4, the ECoG grid did not slide smoothly on the cortex and one column (electrodes 1-3) had to be removed, yielding 4 ECoG electrodes in V1. Two reference wires, common for both microelectrode and ECoG grid were either inserted near the edge of the craniotomy or wounded over the titanium screws on the metal strap which was used to secure the bone on the craniotomy. Other findings based on data recorded from Monkeys 3 and 4 but from a different microelectrode array (implanted in the right hemisphere) have been reported elsewhere^29, 55^.

In case of Monkey 4, we used a hybrid array that had been implanted on a different monkey, but it had to be explanted after 2 days due to complications related to the surgery. One reference wire was lost during the process, and the insulation was removed from the other one (in Monkey 3, insulation from only the tip of the reference wires were removed). This could have led to higher noise in the LFP data collected from Monkey 4 at frequencies above 250 Hz, because the power spectral density appeared to be shallow than other monkeys. It is unlikely that this affected any of the results, since clear gamma rhythm and high-gamma activity were observed in the LFP, which were generally similar to the recordings done earlier using a fresh array implanted in the other hemisphere^29^. Further, ECoG electrodes that were simply placed on the cortex were unaffected by the explantation and showed strong gamma peaks.

All signals were recorded using Blackrock Microsystems data acquisition system (Cerebus Neural Signal Processor). Local field potential (LFP) and multi-unit activity (MUA) were recorded from microelectrode array. LFP and ECoG were obtained by band-pass filtering the raw data between 0.3 Hz (Butterworth filter, first order, analog) and 500 Hz (Butterworth filter, fourth order, digital), sampled at 2 kHz and digitized at 16-bit resolution. MUA was derived by filtering the raw signal between 250 Hz (Butterworth filter, fourth order, digital) and 7,500 Hz (Butterworth filter, third order, analog), followed by an amplitude threshold (set at ∼6.25 (Monkey 1), ∼4.25 (Monkey 2) and ∼5 (Monkeys 3 and 4) of the SDs of the signal).

The data acquisition system has provisions to measure both the impedance of the electrodes as well as potential cross-talk across pairs of electrodes. The similarity in the gamma oscillations recorded in LFP and ECoG signals was not due to potential crosstalk between LFP and ECoG electrodes, which we could measure explicitly. Further, RF centers for LFP and ECoG electrodes were far apart (Figure 5C and 5G), and small stimuli that covered the RF of only one signal produced salient gamma oscillations in that signal but virtually no response in the other, ruling out potential cross-talk influencing our results.

Previously we had also recorded EEG data from Monkeys 3 and 4 simultaneously with the LFP^29^. In this study, EEG signals were found to be extremely noisy. This was because a much larger craniotomy was needed to insert the ECoG array, and consequently a larger titanium mesh, longer plates and more screws were required to secure the bone flap. Further, as this was the second surgery on these monkeys, there was considerable hardware present on the other hemisphere from the first surgery as well. Consequently, there was hardly enough space to put EEG electrodes on the occipital areas, and those signals were noisy.

### Behavioral task

Three separate datasets were used in this study. The first set was used to study the effect of size (‘size study’, Figures 1, 2, 3 and 4) on LFP and ECoG power and were collected from all four monkeys. The second and third data sets were collected from Monkeys 3 and 4 to study the effect of orientation and spatial frequency (‘orientation and spatial frequency study’, Figures 5 and 6) and the effect of contrast (‘contrast study’, Figure 7) on LFP and ECoG power. The behavioral task and stimuli used in these studies are described below in detail.

### Size study

The data set and results from microelectrode recordings from the first two monkeys have been reported previously^17^. The experimental design and behavioral task for ECoG recordings were similar. Monkeys 1 and 2 performed an orientation change task, while two achromatic odd-symmetric stimuli were presented synchronously for 400 ms with an inter-stimulus period of 600 ms. A Grating stimulus of variable size centered on the receptive field of one of the recording electrodes (new location for each session) was presented in the left hemifield for microelectrode recordings and right hemifield for ECoG recordings. The monkeys were cued to attend to a low-contrast Gabor stimulus outside of the receptive field (RF) and respond to a change in the orientation of the Gabor stimulus by 90° in one of the presentations. Monkeys responded by making a saccade within 500 ms of the orientation change. The Gratings were a static stimulus with a spatial frequency of 4 cycles/degree (cpd), full contrast, located at the center of the RF of one of the sites (different recording site each session), one of six different orientations (0°, 30°, 60°, 90°, 120° and 150°) and six different radii (0.3°, 0.72°, 1.14°, 1.56°, 1.98° and 2.4°), chosen pseudo-randomly. For ECoG recordings in Monkey 2, only five radii were presented (up to 1.98°), since the RF center of the ECoG electrode was very fovial (azimuth: 1.16, elevation: 1.83) and the largest stimulus (2.4°) covered the fixation spot. The Gabor stimulus presented outside the RF was also static with an SD of 0.5°, spatial frequency 4 cpd and an average contrast of ∼6% and ∼4.3% for Monkeys 1 and 2. Monkeys 1 and 2 performed the task in 10 and 24 recording sessions for microelectrode recordings (results presented in Ref 17; and 2 and 1 recordings sessions for ECoG recordings (one session for each ECoG electrode).

Monkeys 3 and 4 performed the fixation task while they were in a monkey chair, with their head fixed by the headpost. The monkeys were required to hold their gaze within 2° of a small central dot (0.10° diameter) located at the center of a monitor (BenQ XL2411, LCD, 1280×720 pixels, 100 Hz refresh rate, gamma corrected) and were rewarded with a juice pulse at the end of the trial upon successful fixation. The stimulus was a Grating with a spatial frequency of 4 cpd, full contrast, one of eight different orientations (0°, 22.5°, 45°, 67.5°, 90°, 112.5°, 135° and 157.5°) and six different radii (0.3°, 0.6°, 1.2°, 2.4°, 4.8° and 9.6°), chosen pseudo-randomly, presented for 800 ms with an inter-stimulus period of 700 ms at the RF of one of the recording sites (different recording site each session). The data were collected in 15 (Monkey 3) and 6 (Monkey 4) recording sessions for microelectrode recordings and 5 (Monkey 3) and 4 (Monkey 4) recording sessions for ECoG electrodes.

Only correct trials were used for analysis. For each stimulus size condition, the trials were pooled across orientations to increase the statistical power. The average number of repetitions for each size condition for LFP and ECoG were 182 (range 133 to 288) and 141 (range 129 153) for Monkey 1, 145 (range 106 to 196) and 176 (range 173 to 179) for Monkey 2, 79 (range 37 to 205) and 150 (range 92 to 189) for Monkey 3, and 91 (range 30 to 127) and 115 (range 87 to 153) for Monkey 4.

### Orientation and Spatial frequency tuning study

A full-screen static Grating stimulus was presented for 800 ms with an inter-stimulus period of 700 ms while Monkeys 3 and 4 performed a fixation task. The Gratings were presented at full contrast at one of five spatial frequencies (0.5, 1, 2, 4 and 8 cpd) and one of the eight orientations (0°, 22.5°, 45°, 67.5°, 90°, 112.5°, 135° and 157.5°) chosen pseudo-randomly. The effect of orientation was studied (Figure 5) at spatial frequency which produced highest power in gamma range (4 and 2 cpd for Monkeys 3 and 4). The average number of repetitions for each orientation condition and preferred spatial frequency were 33 (range 28 to 36) for Monkey 3 and 42 (range 37 to 45) for Monkey 4. Similarly, the effect of spatial frequency was studied (Figure 6) at preferred orientation (∼90°) which produced highest gamma power. The average number of repetitions were 33 (range 32 to 36) and 34 (range 15 to 45).

### Contrast study

The stimulus for Monkey 3 was a full-screen Grating at preferred spatial frequency (4 cpd), preferred orientation (90°), one of seven contrasts (100, 50, 25, 12.5, 6.25, 3.125 and 0%) and one of eight different temporal frequencies (tf = 50, 32, 16, 8, 4, 2, 1and 0 cycle per second; counterphase). We studied (Figure 7) the effect of contrast for the static grating (tf = 0 cps); average number of repetitions was 17 (range 16 to 18). For Monkey 4, stimulus was a static full screen Grating at preferred spatial frequency (2 cpd), one of the six contrasts (100, 50, 25, 12.5, 6.25 and 0%) and one of the eight orientations (0°, 22.5°, 45°, 67.5°, 90°, 112.5°, 135° and 157.5°). Contrast tuning was studied at preferred orientation (90°); average number of repetitions was 27 (range 26 to 29). Both monkeys performed a fixation task and stimulus was presented for 800 ms with an inter-stimulus period of 700 ms.

### Electrode selection

Receptive fields were mapped by flashing small Gabor stimuli at various positions on the screen, as described in detail in our previous studies^37, 54^. As in our previous studies, only electrodes for which the RF estimates were stable across days (SD less than 0.1°) were used for further analysis, yielding 27, 71, 77 and 18 microelectrodes and 2, 1, 5 and 4 ECoG electrodes from Monkeys 1, 2, 3 and 4.

For the size study, the smallest stimulus was of radius 0.3°, covering only a few microelectrodes in the visual field. Therefore, for each recording session, we selected electrodes whose RF centers were within 0.2° of the stimulus center. Since we recorded multiple sessions, the same electrode was counted more than once, yielding 56 (24 unique), 141 (66 unique), 62 (40 unique) and 70 (18 unique) electrodes for Monkeys 1-4. Out of this set, we selected electrodes for which the average firing rate was at least 1 spike/s (for an analysis period of 200 to 400 ms for Monkeys 1 and 2 and 250 to 750 ms for Monkeys 3 and 4) for all the stimulus sizes, and a signal-to-noise ratio^56^ greater than 1.5. This yielded 15 (11 unique), 107 (58 unique), 24 (20 unique) and 22 (13 unique) electrodes for further analysis for the four monkeys.

For the orientation, spatial and contrast studies, full screen stimuli were used because that condition produced the strongest gamma. Consequently, firing rates were weak for most sites^29^. Since our primary interest was to compare gamma power, we used the full set of 77 (Monkey 3) and 18 (Monkey 4) microelectrodes and compared the power with 5 (Monkey 3) and 4 (Monkey 4) ECoG electrodes.

### Data analysis

All the data were analyzed using custom codes written in MATLAB (The MathWorks, RRID:SCR_001622). Power spectral density (PSD) and the time-frequency spectra were computed using the multi-taper method with three tapers, implemented in Chronux 2.0 (Bokil et al., 2010, RRID:SCR_005547), an open-source, data analysis toolbox available at http://chronux.org. The baseline period was chosen between −200 to 0 ms for Monkeys 1 and 2 and −500 to 0 ms for Monkeys 3 and 4, where 0 indicates stimulus onset. Stimulus period was chosen between 200 to 400 ms for Monkeys 1 and 2 and 250 to 750 ms for Monkeys 3 and 4 to avoid the stimulus-onset related transients.

Time-frequency difference spectra shown in Figure 2 were obtained by first computing the time-frequency power spectra using a moving window of size 250 ms and a step size of 25 ms and then subtracting the baseline power:

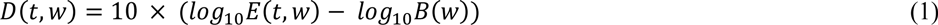

Where *E*(*t, w*) is the mean energy averaged over trials at time t and frequency w, and *B*(*w*) is the baseline energy computed for 500 ms (−500 to 0 ms before stimulus onset). Since subtraction is done on a log scale, this is essentially the log of the ratio of power at any time and the baseline power and has units of decibel (dB). For population data (Figure 2C and 2D), the *D*(*t, w*) values over recording sites were averaged. Note that the baseline energy was calculated across all the stimulus conditions for each recording site.

For the size study, gamma range was chosen between 30 – 65 Hz for all the four monkeys (Figure 3). This was done to accommodate the peak frequency for all stimulus sizes, as gamma peak frequency decreases with an increase in stimulus size^17, 28, 29^. The high-gamma range (150 – 250 Hz) was chosen higher than usual (>80 Hz) to avoid the harmonic of gamma rhythm (∼100 Hz, see Figure 3). The gamma frequency range for orientation and spatial frequency studies, in which a full-screen Grating was presented, was chosen to be 45 – 70 Hz for Monkeys 3 and 4. This was done in congruence with our previous study^29^ which used data from the same two monkeys (but different hemispheres), and to avoid contamination from ‘slow gamma’^29^ which was prominent in Monkey 4. For the contrast study, gamma range was chosen between 20 – 75 Hz. This was done to accommodate peak frequency for all stimulus contrast values, since gamma peak frequency has been to shown to decrease considerably with a reduction in stimulus contrast^27^.

Power in gamma and high-gamma ranges were calculated by first averaging the power values obtained from the PSDs in the corresponding frequency ranges, excluding line noise (60 Hz for Monkeys 1, 2 and 50 Hz for Monkeys 3, 4) and their harmonics. Change in power for each stimulus condition was then calculated as follows:

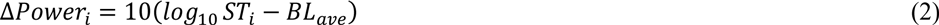

where ST_i_ is the power summed across the frequency range of interest for stimulus condition *i*, and BL_ave_ is the baseline power averaged across conditions (*BL_ave_* = *average*(*log*_10_ *BL_i_*)).

Preferred orientation and orientation selectivity for each recording site were calculated using the following equations:

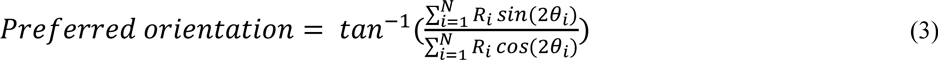

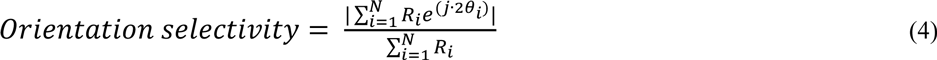

where *θ_i_* and *R_i_* are the orientations and sum of the power in gamma band. N is the total number of orientation values (8).

The slopes (Figure 4) were calculated for stimulus (200 to 400 ms for Monkeys 1, 2 and 250 to 500 ms for Monkeys 3, 4) and baseline (−200 to 0 ms for Monkeys 1, 2 and −500 to 0 ms for Monkeys 3 and 4) periods in high-gamma frequency range (150 – 250 Hz) by fitting the function *log*_10_(*P*) = *m* * *log*_10_(*f*) + *c*, where *P* is the PSD, f is the frequency, *c* is the constant or noise floor and *m* is the slope^40, 57^. In this frequency range, the amplifier roll off is negligible, and therefore the slopes are similar with or without amplifier roll-off correction^40^. We also tested the amplifier noise floor by shorting the inputs and found the power to be at least an order of magnitude lower than the signal power. Therefore, the estimated slopes did not depend on the characteristics of the amplifier.

## Acknowledgements

We thank Dr. John Maunsell for his help in experimental design and data collection from Monkeys 1 and 2 and Steven Sleboda and Vivian Imamura for technical support. We thank Ad-Tech Medical Instrument Corporation and Blackrock Microsystems for the hybrid electrode grid. We also thank Dr. Sebastian Chandu for his assistance in surgeries. This work was supported by Wellcome Trust/DBT India Alliance (500145/Z/09/Z; Intermediate fellowship to SR), Tata Trusts Grant and DBT-IISc Partnership Programme.

## Author contributions

A.D. and S.R. conceptualized the study. S.R. collected the data from Monkeys 1 and 2 and A.D. is responsible for data from Monkeys 3 and 4. A.D. analyzed the data and wrote the first draft of the manuscript. A.D. and S.R. were involved in editing of the manuscript.

## Data availability

The datasets analyzed during the current study are available from the corresponding author on reasonable request.

## References

1. Engel, A. K., Moll, C. K. E., Fried, I. & Ojemann, G. A. Invasive recordings from the human brain: clinical insights and beyond. Nat. Rev. Neurosci. 6, 35–47 (2005).

2. Mukamel, R. et al. Coupling between neuronal firing, field potentials, and FMRI in human auditory cortex. Science 309, 951–954 (2005).

3. Yoshor, D., Bosking, W. H., Ghose, G. M. & Maunsell, J. H. R. Receptive Fields in Human Visual Cortex Mapped with Surface Electrodes. Cereb. Cortex 17, 2293–2302 (2007).

4. Mukamel, R. & Fried, I. Human Intracranial Recordings and Cognitive Neuroscience. Annu. Rev. Psychol. 63, 511–537 (2011).

5. Lachaux, J.-P., Axmacher, N., Mormann, F., Halgren, E. & Crone, N. E. High-frequency neural activity and human cognition: past, present and possible future of intracranial EEG research. Prog. Neurobiol. 98, 279–301 (2012).

6. Morshed, B. I. & Khan, A. A Brief Review of Brain Signal Monitoring Technologies for BCI Applications: Challenges and Prospects. J. Bioeng. Biomed. Sci. 4, 1–10 (2014).

7. Yang, T., Hakimian, S. & Schwartz, T. H. Intraoperative ElectroCorticoGraphy (ECog): indications, techniques, and utility in epilepsy surgery. Epileptic. Disord. 271–279 (2014). doi:10.1684/epd.2014.0675

8. Winawer, J. & Parvizi, J. Linking Electrical Stimulation of Human Primary Visual Cortex, Size of Affected Cortical Area, Neuronal Responses, and Subjective Experience. Neuron 92, 1213–1219 (2016).

9. Kucewicz, M. T. et al. Dissecting gamma frequency activity during human memory processing. Brain J. Neurol. 140, 1337–1350 (2017).

10. Parvizi, J. & Kastner, S. Promises and limitations of human intracranial electroencephalography. Nat. Neurosci. 21, 474–483 (2018).

11. Crone, N. E., Sinai, A. & Korzeniewska, A. High-frequency gamma oscillations and human brain mapping with electrocorticography. Prog. Brain Res. 159, 275–295 (2006).

12. Crone, N. E., Korzeniewska, A. & Franaszczuk, P. J. Cortical γ responses: searching high and low. Int. J. Psychophysiol. Off. J. Int. Organ. Psychophysiol. 79, 9–15 (2011).

13. Ray, S., Crone, N. E., Niebur, E., Franaszczuk, P. J. & Hsiao, S. S. Neural Correlates of High-Gamma Oscillations (60–200 Hz) in Macaque Local Field Potentials and Their Potential Implications in Electrocorticography. J. Neurosci. 28, 11526–11536 (2008).

14. Liu, J. & Newsome, W. T. Local field potential in cortical area MT: stimulus tuning and behavioral correlations. J. Neurosci. Off. J. Soc. Neurosci. 26, 7779–7790 (2006).

15. Manning, J. R., Jacobs, J., Fried, I. & Kahana, M. J. Broadband shifts in local field potential power spectra are correlated with single-neuron spiking in humans. J. Neurosci. Off. J. Soc. Neurosci. 29, 13613–13620 (2009).

16. Jia, X., Smith, M. A. & Kohn, A. Stimulus selectivity and spatial coherence of gamma components of the local field potential. J. Neurosci. Off. J. Soc. Neurosci. 31, 9390–9403 (2011).

17. Ray, S. & Maunsell, J. H. R. Different Origins of Gamma Rhythm and High-Gamma Activity in Macaque Visual Cortex. PLoS Biol 9, e1000610 (2011).

18. Rodriguez, E. et al. Perception’s shadow: long-distance synchronization of human brain activity. Nature 397, 430–433 (1999).

19. Fries, P., Reynolds, J. H., Rorie, A. E. & Desimone, R. Modulation of Oscillatory Neuronal Synchronization by Selective Visual Attention. Science 291, 1560–1563 (2001).

20. Pesaran, B., Pezaris, J. S., Sahani, M., Mitra, P. P. & Andersen, R. A. Temporal structure in neuronal activity during working memory in macaque parietal cortex. Nat. Neurosci. 5, 805–811 (2002).

21. Womelsdorf, T., Fries, P., Mitra, P. P. & Desimone, R. Gamma-band synchronization in visual cortex predicts speed of change detection. Nature 439, 733–736 (2006).

22. Melloni, L. et al. Synchronization of neural activity across cortical areas correlates with conscious perception. J. Neurosci. Off. J. Soc. Neurosci. 27, 2858–2865 (2007).

23. Colgin, L. L. et al. Frequency of gamma oscillations routes flow of information in the hippocampus. Nature 462, 353–357 (2009).

24. Gregoriou, G. G., Gotts, S. J., Zhou, H. & Desimone, R. High-frequency, long-range coupling between prefrontal and visual cortex during attention. Science 324, 1207–1210 (2009).

25. Henrie, J. A. & Shapley, R. LFP Power Spectra in V1 Cortex: The Graded Effect of Stimulus Contrast. J. Neurophysiol. 94, 479–490 (2005).

26. Gieselmann, M. A. & Thiele, A. Comparison of spatial integration and surround suppression characteristics in spiking activity and the local field potential in macaque V1. Eur. J. Neurosci. 28, 447–459 (2008).

27. Ray, S. & Maunsell, J. H. R. Differences in Gamma Frequencies across Visual Cortex Restrict Their Possible Use in Computation. Neuron 67, 885–896 (2010).

28. Jia, X., Xing, D. & Kohn, A. No Consistent Relationship between Gamma Power and Peak Frequency in Macaque Primary Visual Cortex. J. Neurosci. 33, 17–25 (2013).

29. Murty, D. V. P. S., Shirhatti, V., Ravishankar, P. & Ray, S. Large visual stimuli induce two distinct gamma oscillations in primate visual cortex. J. Neurosci. 2270–17 (2018). doi:10.1523/JNEUROSCI.2270-17.2017

30. Rols, G., Tallon-Baudry, C., Girard, P., Bertrand, O. & Bullier, J. Cortical mapping of gamma oscillations in areas V1 and V4 of the macaque monkey. Vis. Neurosci. 18, 527–540 (2001).

31. Adjamian, P. et al. Induced visual illusions and gamma oscillations in human primary visual cortex. Eur. J. Neurosci. 20, 587–592 (2004).

32. Hall, S. D. et al. The missing link: analogous human and primate cortical gamma oscillations. NeuroImage 26, 13–17 (2005).

33. Swettenham, J. B., Muthukumaraswamy, S. D. & Singh, K. D. Spectral Properties of Induced and Evoked Gamma Oscillations in Human Early Visual Cortex to Moving and Stationary Stimuli. J. Neurophysiol. 102, 1241–1253 (2009).

34. Muthukumaraswamy, S. D. & Singh, K. D. Visual gamma oscillations: the effects of stimulus type, visual field coverage and stimulus motion on MEG and EEG recordings. NeuroImage 69, 223–230 (2013).

35. Perry, G., Hamandi, K., Brindley, L. M., Muthukumaraswamy, S. D. & Singh, K. D. The properties of induced gamma oscillations in human visual cortex show individual variability in their dependence on stimulus size. NeuroImage 68, 83–92 (2013).

36. Hermes, D., Miller, K. J., Wandell, B. A. & Winawer, J. Stimulus Dependence of Gamma Oscillations in Human Visual Cortex. Cereb. Cortex N. Y. N 1991 25, 2951–2959 (2015).

37. Dubey, A. & Ray, S. Cortical Electrocorticogram (ECoG) Is a Local Signal. J. Neurosci. 39, 4299–4311 (2019).

38. Mitra, P. P. & Pesaran, B. Analysis of dynamic brain imaging data. Biophys. J. 76, 691–708 (1999).

39. Bokil, H., Andrews, P., Kulkarni, J. E., Mehta, S. & Mitra, P. Chronux: A Platform for Analyzing Neural Signals. J. Neurosci. Methods 192, 146–151 (2010).

40. Shirhatti, V., Borthakur, A. & Ray, S. Effect of Reference Scheme on Power and Phase of the Local Field Potential. Neural Comput. 28, 882–913 (2016).

41. Hubel, D. H. & Wiesel, T. N. Receptive fields, binocular interaction and functional architecture in the cat’s visual cortex. J. Physiol. 160, 106 (1962).

42. Hubel, D. H. & Wiesel, T. N. Receptive fields and functional architecture of monkey striate cortex. J. Physiol. 195, 215 (1968).

43. Hubel, D. H. & Wiesel, T. N. Ferrier Lecture: Functional Architecture of Macaque Monkey Visual Cortex. Proc. R. Soc. Lond. B Biol. Sci. 198, 1–59 (1977).

44. Berens, P., Keliris, G. A., Ecker, A. S., Logothetis, N. K. & Tolias, A. S. Comparing the Feature Selectivity of the Gamma-Band of the Local Field Potential and the Underlying Spiking Activity in Primate Visual Cortex. Front. Syst. Neurosci. 2, (2008).

45. Xing, D., Yeh, C.-I. & Shapley, R. M. Spatial Spread of the Local Field Potential and its Laminar Variation in Visual Cortex. J. Neurosci. 29, 11540–11549 (2009).

46. Canolty, R. T. et al. High gamma power is phase-locked to theta oscillations in human neocortex. Science 313, 1626–1628 (2006).

47. Buschman, T. J. & Miller, E. K. Top-down versus bottom-up control of attention in the prefrontal and posterior parietal cortices. Science 315, 1860–1862 (2007).

48. He, B. J., Zempel, J. M., Snyder, A. Z. & Raichle, M. E. The temporal structures and functional significance of scale-free brain activity. Neuron 66, 353–369 (2010).

49. Pesaran, B. et al. Investigating large-scale brain dynamics using field potential recordings: analysis and interpretation. Nat. Neurosci. 21, 903–919 (2018).

50. Srinath, R. & Ray, S. Effect of amplitude correlations on coherence in the local field potential. J. Neurophysiol. 112, 741–751 (2014).

51. Xu, X., Anderson, T. J. & Casagrande, V. A. How do functional maps in primary visual cortex vary with eccentricity? J. Comp. Neurol. 501, 741–755 (2007).

52. Uhlhaas, P. J. & Singer, W. Abnormal neural oscillations and synchrony in schizophrenia. Nat. Rev. Neurosci. 11, 100–113 (2010).

53. Uhlhaas, P. J. & Singer, W. Neuronal dynamics and neuropsychiatric disorders: toward a translational paradigm for dysfunctional large-scale networks. Neuron 75, 963–980 (2012).

54. Dubey, A. & Ray, S. Spatial spread of local field potential is band-pass in the primary visual cortex. J. Neurophysiol. 116, 1986–1999 (2016).

55. Shirhatti, V. & Ray, S. Long-wavelength (reddish) hues induce unusually large gamma oscillations in the primate primary visual cortex. Proc. Natl. Acad. Sci. U. S. A. 115, 4489–4494 (2018).

56. Kelly, R. C. et al. Comparison of Recordings from Microelectrode Arrays and Single Electrodes in the Visual Cortex. J. Neurosci. 27, 261–264 (2007).

57. Miller, K. J., Sorensen, L. B., Ojemann, J. G. & Nijs, M. den. Power-Law Scaling in the Brain Surface Electric Potential. PLOS Comput. Biol. 5, e1000609 (2009).

